# Super Resolved Structural Imaging of Mitochondrial Network using Orange Emissive Carbon Nanodot as Fluorescent Probe

**DOI:** 10.1101/2024.12.13.628462

**Authors:** Richa Garg, Runmi Kundu, Kush Kaushik, Abdul Salam, Chayan Kanti Nandi

## Abstract

Super-resolution microscopy (SRM), coupled with appropriate bright and photostable fluorescent probes, has revolutionized the ability to study organelle dynamics with unprecedented spatial and temporal resolution. An increasing trend of designing nanomaterial probes that has unprecedented advantages over the organic molecular probes has become the frontier in SRM based imaging of subcellular organelles. Herein, we report the development of orange-emissive fluorescent carbon nanodots (CNDs) via a one-pot synthesis that has excellent capabilities to target mitochondria. Spectroscopic analysis confirms the presence of guanidine on the surface of CNDs, thus facilitating its ability to selectively target mitochondria. The CNDs were highly capable for the super resolution radial fluctuation (SRRF) imaging of mitochondrial network and the morphology. The synthesized CNDs exhibited excellent photoluminescent properties along with high biocompatibility and non-toxicity, which could be used for their application in mitochondria-based imaging modalities.

## Introduction

Fluorescent imaging and a detailed understanding of interconnected mitochondria networks with nanometer scale resolution promises to reveal their ultrastructural dynamics, which is essential for countering physiological disorders. Super-resolution microscopy (SRM) plays a significant role in the comprehensive analysis of cellular cytoskeletal dynamics such as lysosomes, mitochondria, Golgi body and ER in live cells^[1]^. SRM is a nanoscopic technique that helps to visualize the organelle structure and their dynamics within the cell at a nanoscale level, hence helping to uncover the ultrastructural dynamics of tiny organelles^[2–4]^. A bright, non-toxic and biocompatible fluorescent probe is often required to get the high quality SRM image. In recent years, remarkable progress has been made in developing fluorescent probes such as quantum dots, gold nanoclusters, metal complexes, and polymer dots, which are specific for particular organelles ^[5–7]^. Despite this technological advancement, the complex structure of organellar morphology still faces several challenges in accurately identifying and analyzing data in a complex biological environment.

Mitochondria is an essential organelle within a cell that sustains cellular energy by generating ATP^[8]^. Mitochondria exist as a dynamic and interconnected network due to continuous fusion and fission mitochondrial events within the cells that finally help in cellular homeostasis^[9]^. Despite advances in cellular biology that have improved our knowledge of mitochondrial structure, many unknowns still hinder efforts to solve mitochondrial-related problems. So, it is essential to thoroughly map and understand mitochondrial morphology. Therefore, various advanced imaging modalities have been utilized, significantly contributing to a deeper understanding of mitochondrial morphology^[10–12]^. Out of several, SRM found to be an effective technique to directly visualize the cellular organelle and also their dynamics down to the nanometer level in a living cell for a longer period of time without affecting the cellular environment^[13–16]^.

The SRM technique desired efficient and suitable fluorescent probes with appropriate photoluminescent properties such as high brightness and long-lived in biological milieu without producing much toxicity^[17]^. Although, over the years, many synthetic organic dyes^[6]^, fluorescent proteins^[7]^, and quantum dots^[5]^ were used for SRM imaging, they suffer from low photostability, low brightness, rapid photobleaching, toxic effect^[18]^, that limits the applications in SRM imaging. These issues impede their effectiveness in real-time monitoring and long-term tracking applications. Consequently, recent research has focused on developing analogous nanomaterial-based fluorophores to address these limitations and improve long-term organellar tracking efficiency. Among these, CNDs are promising fluorescent nanomaterials due to their strong biocompatibility, excellent cell permeability, bright fluorescence, high photostability, and simple synthesis. These characteristics make CNDs compelling candidates for bioimaging applications^[19–22]^.

Current mitochondria-targeting approaches exploit their high negative potential and hydrophobic double membrane structure. Essential design features for these probes include a cationic charge for electrostatic binding and lipophilicity for interaction with the hydrophobic membrane. Triphenylphosphonium (TPP), dequalinium, guanidine salt, and rhodamine are commonly used for effective mitochondria targeting^[23–27]^. For example, Wu et al. developed a mitochondrial specific nanoprobe by attaching triphenylphosphonium (TPP) to CNDs synthesized from o-phenylenediamine, which exhibited an emission wavelength of 575 nm. Ray et al. developed a nanoprobe with a green-emissive quantum dot (QD) or γ-Fe_2_O_3_ core and a polyacrylate shell functionalized with arginine and primary amine groups for enhanced mitochondrial targeting^[28]^. Hazra et al. developed spherical micelle-type nanostructures using an amphiphile and adorned them with green-emitting CNDs derived from 2,3-dichloro-5,6-dicyano-1,4-benzoquinone (DDQ), effectively imaging mitochondria^[29]^. However, developing mitochondria-specific CNDs in a single-step synthesis method with longer wavelength emission remains challenging, especially for SRM imaging.

In this study, we have synthesized mitochondria-specific CNDs using a one-step method with o-phenylenediamine (OPDA) and guanidine hydrochloride as precursors. Several chemical characterization techniques confirmed Guanidine presence on the CNDs surface, facilitating its ability to target mitochondria specifically. Confocal microscopy images also demonstrate effective mitochondrial targeting in mammalian cells. Based on better biocompatibility and mitochondria specificity, the developed small size CNDs showed an excellent probe for the SRM imaging of mitochondrial networks inside cellular environment. CNDs were highly capable of capturing the images of mitochondria with an average width of 0.5 µM and an average length of 3.5 µM, which are well matched with the reported values.

## Result and Discussion

The synthesis of CNDs was achieved through a bottom-up solvothermal treatment method using o-phenylenediamine (OPDA) and guanidine hydrochloride as the precursor materials. Typically, 108 mg of OPDA and 191 mg of guanidine hydrochloride were mixed. Subsequently, 4.5 mL of distilled water and hydrochloric acid were added to the vial with a continuous stirring to ensure homogeneity. Next, it was carefully transferred into a Teflon-lined 25 mL stainless hydrothermal container, which was then placed inside an oven preheated to 180 °C. The reaction was continued for 12 hours. After 12 hours, the container was removed from the oven and allowed to cool gradually to room temperature, as shown in **Figure 1a**. The obtained reaction product was then centrifuged at 7000 rpm for 30 minutes to remove unwanted impurities. After that, dialysis was performed to get a pure sample of CNDs. The brown solid CNDs were then obtained by freeze-drying.

**Figure 1:**
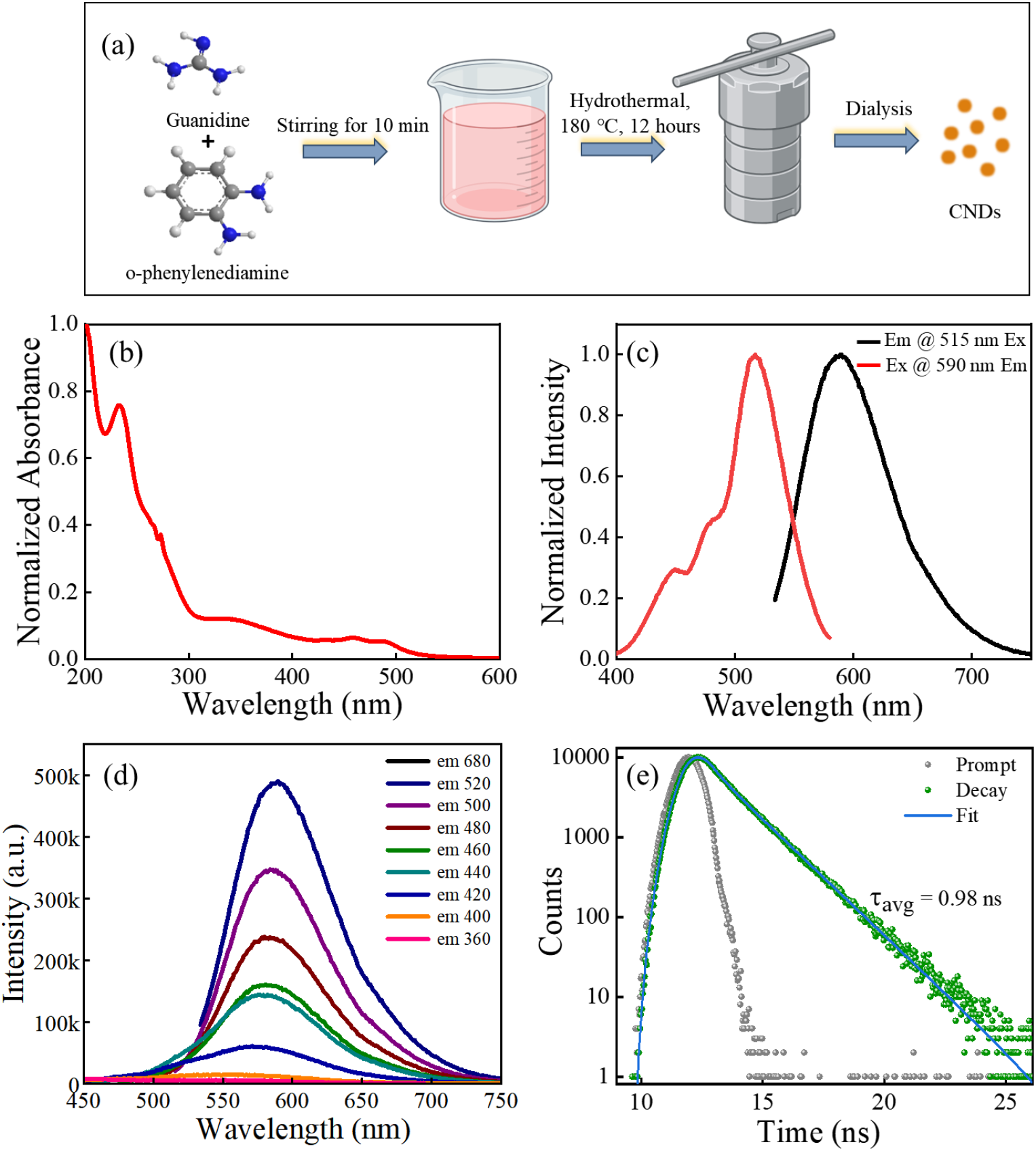
(a) Synthetic protocol of CNDs. (b) Normalized UV–vis spectrum of CNDs. (c) Normalized excitation and emission spectra of CNDs. (d) Full emission spectra of CNDs excited at different wavelengths. (e) Lifetime decay curve of CNDs.

Both chemical and optical characterization were carried out to determine the nature of CNDs. The optical properties of the synthesized CNDs were examined by analyzing their UV-Vis absorption, steady-state fluorescence, and time-resolved fluorescence spectra after dispersing them in deionized water. The normalized absorption **(Figure 1b)** peaks below 400 nm are attributed to surface states transitions related to n → π* transitions in C=N bonds and π → π* transition in aromatic sp^2^ carbons (aromatic C=C bonds) within the carbon cores. Peaks above 400 nm are due to molecular state transitions, as was proposed earlier^[22]^. **Figure 1c** shows a normalized emission spectrum with a peak maximum at 590 nm when excited at 515 nm. While the excitation spectrum with a peak maximum at 515 nm was obtained when the emission was fixed at 590 nm. The detailed emission fluorescence spectra of CNDs **(Figure 1d)** with different excitation wavelengths were also captured, showing an emission peak around 590 nm. This behaviour is attributed to the presence of surface defects or molecular states within the CNDs structure, which act as the dominant emission centers. We also measured the fluorescence lifetime of CNDs **(Figure 1e)**, which was found to be 0.98 ns with an excitation at 454 nm.

The graphitization was confirmed through raman spectra of CNDs **(Figure 2a)**. The typical peaks are located approximately at 1362 cm^-1^ (D-band) and 1570 cm^-1^ (G-band), corresponding to the disorders, structural defects, and characterizing the ordered in-plane vibrations of sp^2^ graphitic carbon respectively^[30]^. Along with D and G bands, raman spectra also show N-H bending vibrations at 1254 cm^-1^ and C=N, C=C stretching vibrations at 1475 cm^-1^ and 1627 cm^-1,^ respectively^[31]^. The raman spectra indeed proved graphitic core structure in CNDs. Thermogravimetric analysis (TGA) was conducted to assess the sample’s mass loss upon heating, providing insights into CNDs thermal stability. **Figure 2b** illustrates a mass loss between 25 to 220 °C, followed by a significant mass loss until reaching 380 °C. The substantial mass loss at lower temperatures (up to 380 °C) indicates the presence of surface-state molecules, which are less thermally stable and volatile. Conversely, the species remaining stable up to 800 °C, could be attributed to the formation of thermally stable aggregated cores^[32]^. Transmission electron microscope (TEM) analysis of CNDs confirmed the dot-like morphology in **Figure 2c** with an average size of ∼ 3.0 nm **(Figure 2d)**. The height profiling of CNDs was done by atomic force microscopy (AFM), shown in **Figure 2e**. The data reveals the topological height to be 4 nm, confirming the stacking of 10-12 graphene-like sheets. The X-ray diffraction pattern of CNDs displayed multiple sharp peaks rather than a broad peak, indicating that the CNDs are polycrystalline, as illustrated in **Figure 2f**.

**Figure 2:**
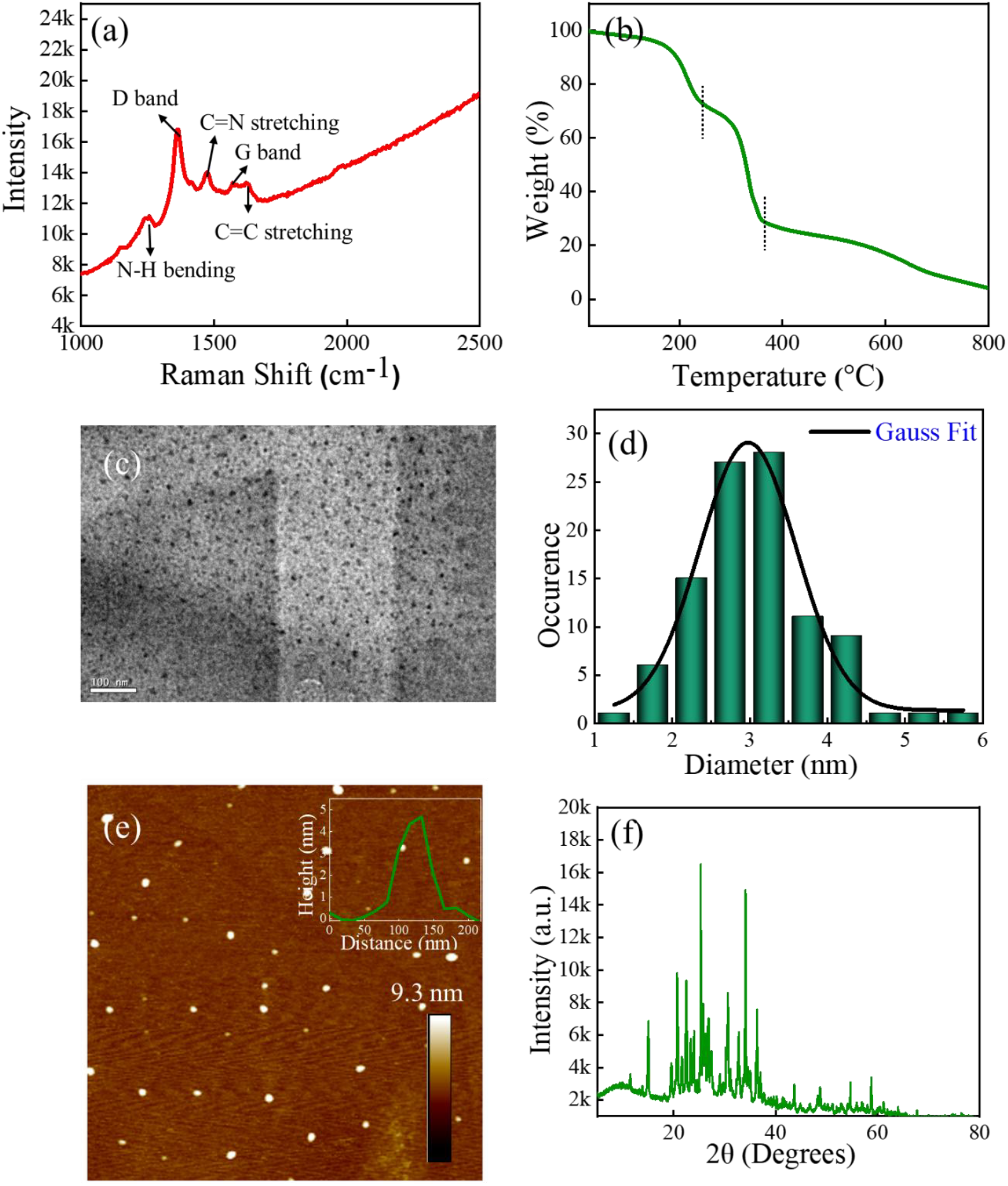
Characterization of CNDs. (a) Raman spectra of CNDs upon excitation with 532 nm laser. The D & G bands were appeared at 1364 cm-1 (D-band) and 1570 cm-1 (G-band) along with other C=C and C=N stretching and N-H bending vibrations over the fluorescence envelop. (b) Thermogravimetric analysis of CNDs. (c) TEM image of CNDs (Scale bar 100 nm). (d) Statistical analysis of CNDs size of approximately 200 particles. (e) AFM image of CNDs. Inset is showing an average height profile of ∼4 nm. (f) PXRD spectra of CNDs.

Figure 3 depicts a comparison of the FTIR spectra of OPDA, guanidine, and CNDs. X-ray photoelectron spectroscopy (XPS) analysis was performed to determine the chemical composition and surface functional groups of CNDs. The Fourier transform infrared (FTIR) spectra indicate the characteristic peaks of CNDs at 3408 cm^-1^ (N-H stretching), 3147 cm^-1^ (C-H stretching), and 1622 cm^-1^ (C=C stretching) (**Figure 3a**). These findings suggest that the surface of the CNDs is enriched with N-H groups. The survey revealed **(Figure 3b)** that CNDs mainly contain carbon (C), nitrogen (N), oxygen (O), and chlorine (Cl), with their respective contents being 58.02%, 27.21%, 5.58%, and 9.18%. The high-resolution N1s spectrum **(Figure 3c)** displayed a peak at 400.3 eV, indicating the presence of -NH-, where the peak at 398.9 eV was attributed to -NH_2_. The O1s spectrum **(Figure 3d)** at 533.6, 532.5 and 531.1 eV confirmed due to C-O, O=C-O, and C=O groups. The C1s high-resolution spectra **(Figure 3e)** show binding energies at 284.8 eV for C−C/C=C, 285.8 eV for C−N, and 289.5 eV for C=O. Additionally, the Cl2p spectrum showed **(Figure 3f)** peaks at 198.1 and 199.6 eV, representing the C-Cl bond with two different spin states. The hypothesized structure of CNDs is also shown in **Scheme 1a**. Both XPS and FTIR analyses support the presence of -NH_2_ groups on the surface, indicating a significant presence of C-NH_2_ groups that facilitate targeting mitochondria, as shown in **Scheme 1b**.

**Figure 3:**
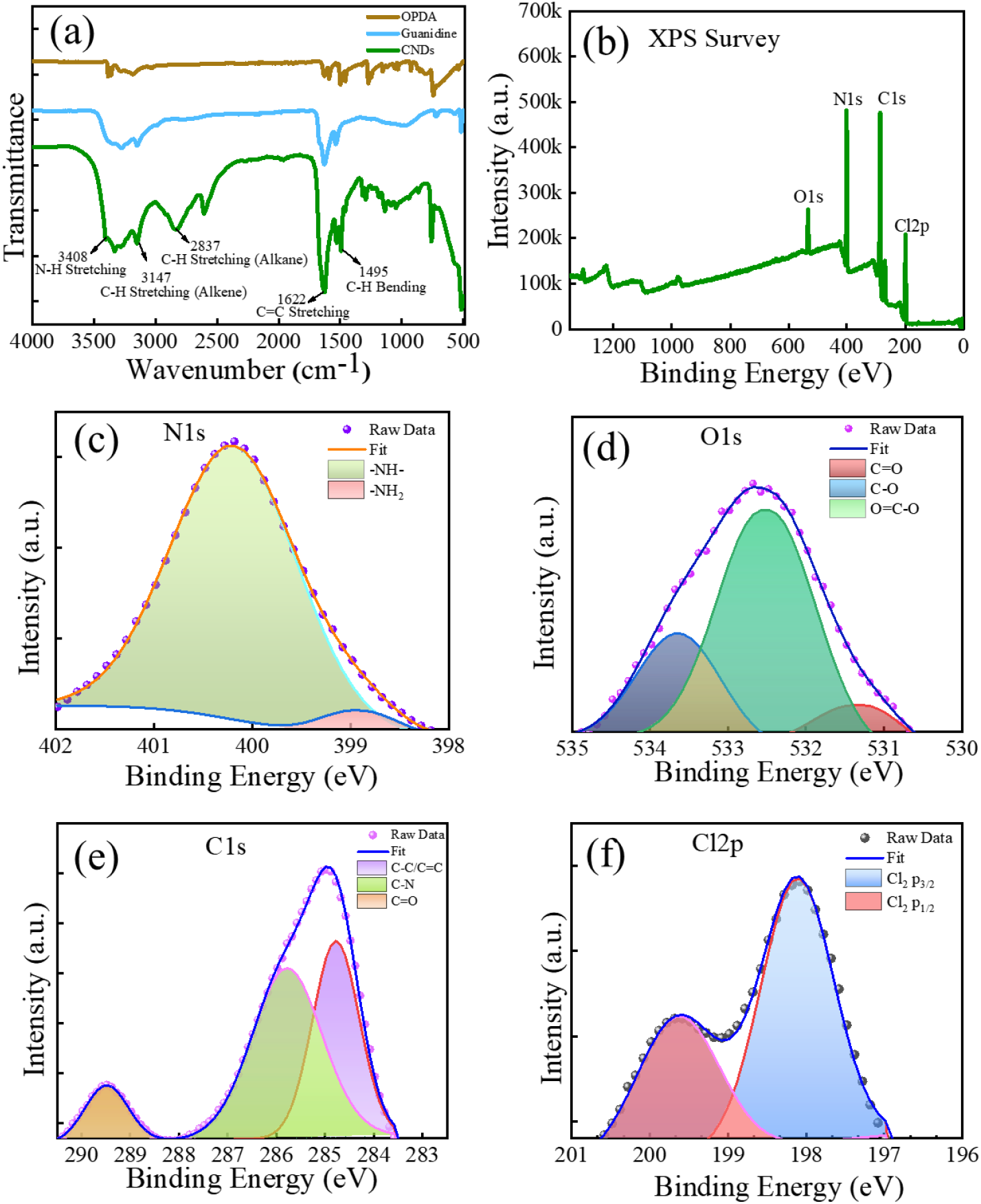
(a) FT-IR spectra of the CNDs (green) and the Guanidine (blue) and OPDA (Brown). XPS survey of CNDs. The high-resolution XPS peaks of (c) N Is, (d) O 1s and (e) C 1s, and Cl 2p, respectively.

**Scheme 1.**
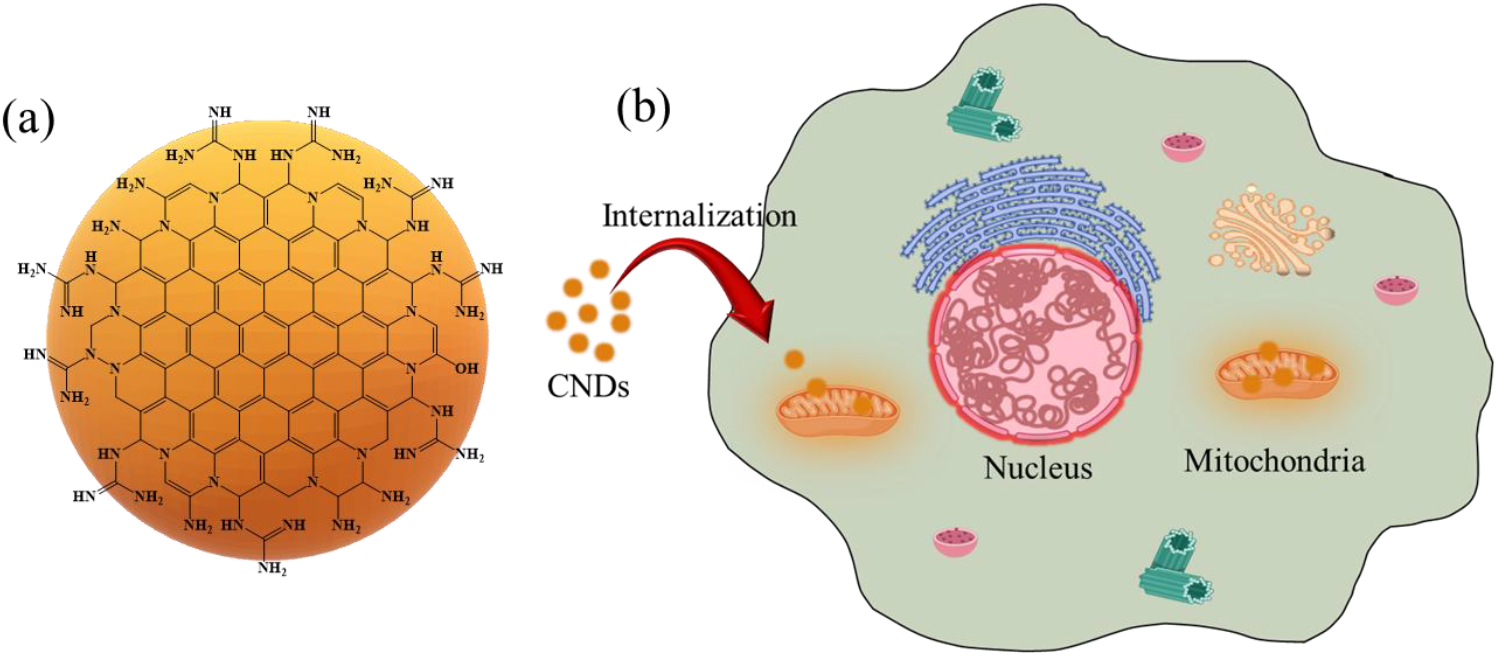
. (a) Schematic of the proposed structure of CNDs (b) CNDs applications as a fluorescent nano dot for specific mitochondrial imaging.

Before employing CNDs for mitochondria bioimaging, the XTT assay was evaluated inHeLa cells. XTT assay showed **(Figure 4)** that over 80% of HeLa cells survived even after 6, 12, and 24 h incubation, indicating the excellent cell viability of CNDs. Based on the excellent optical and biocompatibility properties of CNDs, we performed cellular bioimaging to validate the mitochondrial specificity of CNDs in non-cancerous (HEK) and cancerous (HeLa) cells by utilizing confocal microscopy. CNDs and MitoTracker Green (MTG) were incubated in HEK and HeLa cells for 1 hour, followed by washing to remove any unbound CNDs. The images were captured using 561 nm and 488 nm lasers to excite the CNDs and MitoTracker Green, respectively.

**Figure 4:**
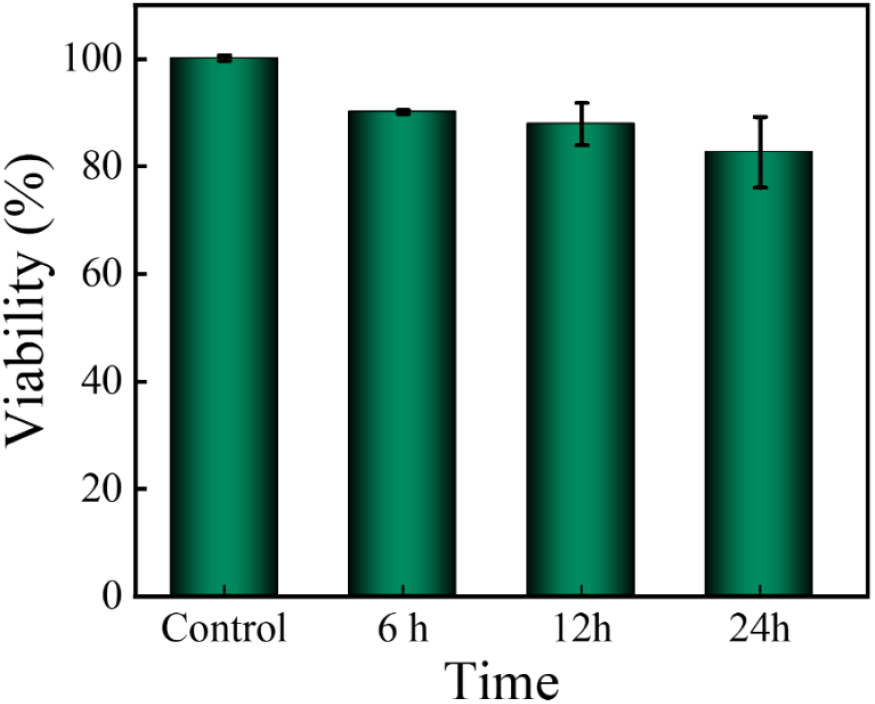
XTT results of HeLa cell viability after coincubation with CNDs for 6, 12, and 24 h at the concentrations of 85 μg/mL.

Figure 5 shows the confocal images where the green color represents MitoTracker Green, while the yellow color indicates the CNDs. The yellow fluorescence of CNDs was consistent with green fluorescence in the merged image, showing their excellent colocalization with mitochondria throughout the cells. The degree of colocalization was quantified using ImageJ software, yielding a Pearson correlation coefficient greater than 0.8 in both HEK and HeLa cells, which confirms that the CNDs specifically target mitochondria in both cancerous and non-cancerous cells. These results demonstrate that the CNDs offer valuable insights into mitochondrial morphology, function, and activity in various cell types. The guanidine moieties on the surface of the CNDs enhance their ability to recognize and effectively label mitochondria^[24–26]^. Consequently, these newly synthesized CNDs can serve as a reliable mitochondrial labeling probe for both cancerous and non-cancerous cells.

**Figure 5:**
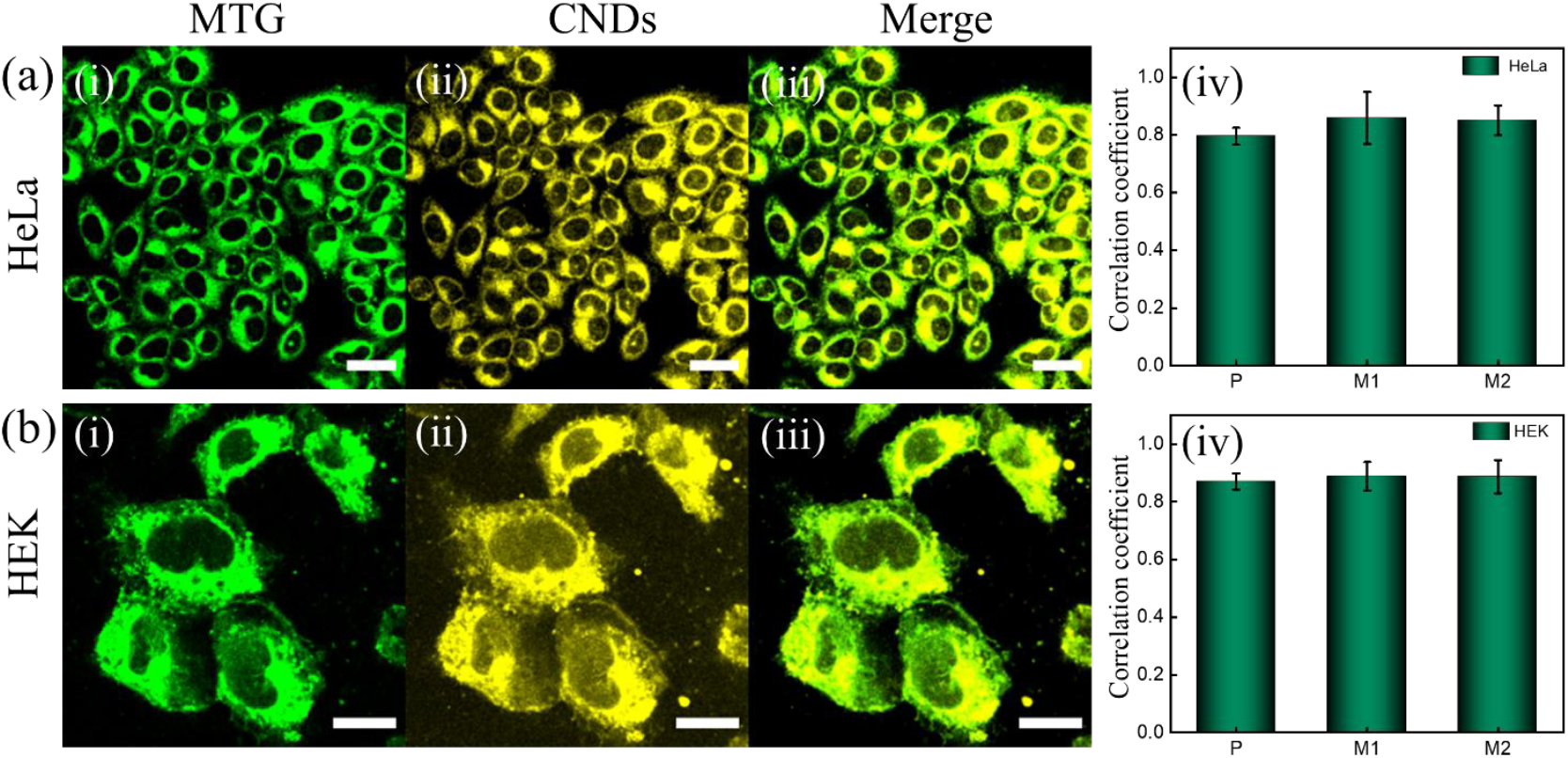
(a) Confocal fluorescence images for HeLa cells co-stained with Mito-tracker and CNDs. The cells were excited at 488 nm and 560 nm, respectively. i & ii represents the fluorescence images of MitoTracker and CNDs respectively (scale bar 30 µm). iii represent merged images of i & ii. (b) Confocal fluorescence images for HEK cells co-stained with Mito-tracker and CNDs. Here i & ii represents the fluorescence images of MitoTracker and CNDs respectively (scale bar 20 µm). iii represent merged images of i & ii. iv represent colocalization of MitoTracker Green (MTG) and CNDs in HeLa, and HEK cells. The Pearson correlation coefficient (P) and Manders’ overlap coefficients (M1 and M2) value of greater than 0.8 confirms the colocalization.

Mitochondria continuously change their shape, size and undergo fission and fusion reactions to maintain cellular homeostasis in normal conditions. After confirming the specificity of CNDs towards the mitochondria, we performed the SRRF to obtain high-resolution images of the interconnected mitochondrial network in HeLa cells. **Figure 6** shows the transmission detection image (TD) and their corresponding SRRF image of the mitochondrial network in HeLa cells stained with CNDs. It shows long thread-like structures of an interconnected mitochondrial network. We also quantitatively analyzed mitochondrial dimensions such as width and length in HeLa cells under normal conditions^[33]^. For this, we analyzed five cells and examined over 100 mitochondria from the entire cytoplasm to minimize biases. Under normal conditions, the average length and width of mitochondria in HeLa cells were ∼0.5 µm and ∼3.5 µm, respectively. These results suggested that the CNDs reported in this work are highly capable to specifically stain mitochondria and can capture the super-resolved mitochondrial network.

**Figure 6:**
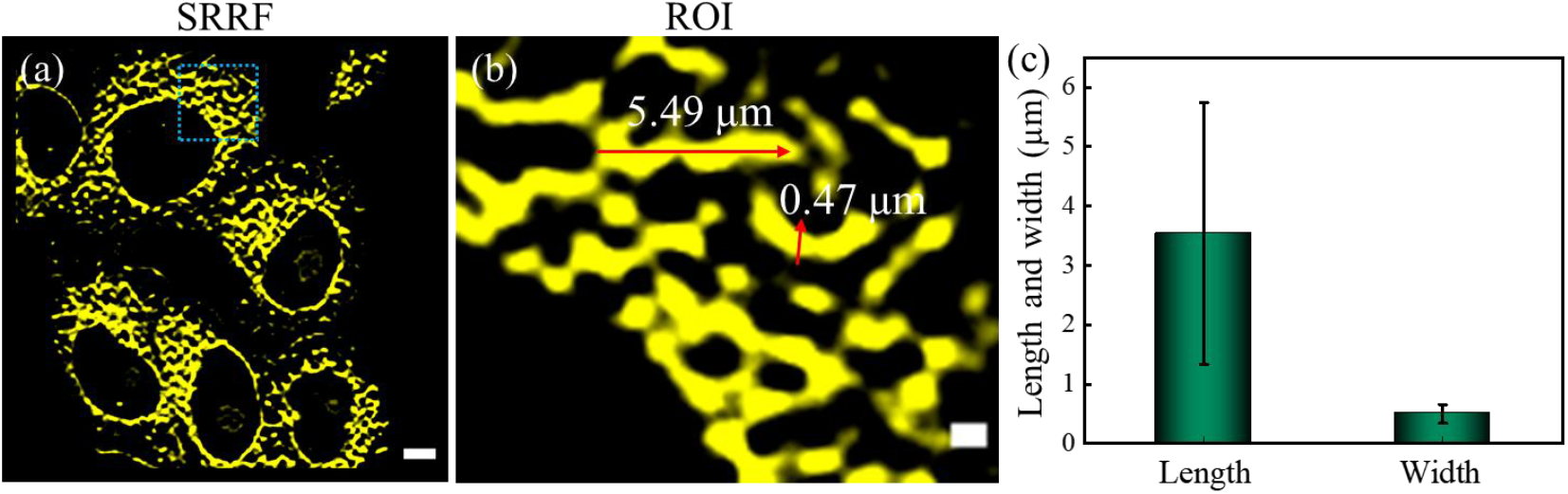
(a) SRRF image of mitochondrial network stained with CNDs in HeLa cells (scale bar 5 µm). b represents the zoomed image of marked area of a (scale bar 1 µm). (c) Quantitative length and width analysis of mitochondria in HeLa cells under normal conditions.

## Conclusions

In summary, we have developed a new highly efficient, biocompatible, mitochondria-specific orange-emissive CNDs using OPDA and guanidine through a one-pot method, eliminating the need for additional conjugation with targeting or recognition molecules. By applying our probes in SRM imaging, we successfully recorded the super-resolved interconnected mitochondrial Network in HeLa cells. The average width and length of mitochondria in Hela cells under normal conditions were ∼0.5 µm and ∼3.5 µm, respectively. Spectroscopic analysis confirms the presence of guanidine on the CNDs surface, facilitating its ability to target mitochondria specifically. Confocal imaging demonstrates effective mitochondrial targeting in mammalian cells, as verified by colocalization with MitoTracker. These CNDs are promising new nanomaterials for advancing targeted drug delivery and monitoring mitochondrial dynamics at the nanoscale during normal and disease conditions.

## Data availability

All experimental and characterization data that support this article are available in the main manuscript and its ESI.

## Supporting information

Electronic Supplementary Information

## Acknowledgments

CKN acknowledges to the Science & Engineering Research Board (Project No. IITM/SERB/CKN/310) for the financial support. CKN is also acknowledges funding form the Council of Scientific & Industrial Research (IITM/CSIR/CKN/449). CKN thanks IIT Mandi for providing a state-of-the-art instrumentation facility including the cell culture. The authors thank the Advanced Materials Research Centre (AMRC), IIT Mandi for experimental facilities. KK and AS thank the Ministry of Education (MoE), India, for the research scholarship. RG acknowledges the Prime Minister Research Fellowship (PMRF).

## Author contributions

RG and RK synthesized and performed all the photophysical characterizations. RG performed all cell culture experiments with the help of AS. KK helped in SRRF imaging and analysis. RG wrote the article with the help of RK and CKN. CKN mentored and guided the whole project.

