## Supplementary material for "Super Resolved Structural Imaging of Mitochondrial Network using Orange Emissive Carbon Nanodot as Fluorescent Probe": Electronic Supplementary Information

### Materials and Methods

#### Chemicals and Materials:

O-phenylenediamine (OPDA) and Guanidine both chemicals were purchased from Sigma Aldrich. HCl was purchased from rankem. Dulbecco's Modified Eagle Medium (DMEM), 1% Antibiotic-Antimicotic, Penstrap, and fetal bovine serum (FBS) were purchased from Gibco. All chemicals were of analytical quality and did not require further purifications. All glassware and hydrothermal were washed with aqua regia ($3HCl:1HNO_3$), followed by washing with ethanol, acetone, and rinsed several times with double distilled water. Double-distilled (18.3 MΩ) deionized water (Elga Purelab Ultra, Vivendi water system Ltd, India) was used throughout the whole experimental process. The cell lines used in this study were purchased from the National Centre for Cell Science (NCCS) Complex, Pune, Maharashtra, India - 411 007.

#### Characterization Techniques:

#### UV-Visible spectroscopy

The UV-Vis of CNDs was measured on a (50 W halogen lamp (2000 h life)) Shimadzu UV-Vis 2450 spectrophotometer. CNDs were placed in a transparent quartz cuvette with 1 ml volume and 10 mm path length for absorption spectra.

#### Steady-state fluorescence spectroscopy

Steady-state fluorescence of CNDs was measured using a Horiba Fluorolog-3 spectrofluorometer. The fluorescence spectra were measured using a 1 ml quartz cuvette having 10 mm path length. All the measurements were repeated at least two times.

#### Fluorescence lifetime

The fluorescence lifetime was examined using the Horiba scientific Delta Flex TCSPC system with 574 nm Pulsed LED Sources. Ludox has been used as an IRF. The fluorescence lifetime was identified by bi-exponentially fitting, the photon decays in different channels with a chi-squared value < 1.1.

**Thermogravimetric Analysis (TGA)**

Thermal properties were measured by Perkin Elmer Pyris Thermogravimetric analyzer from 25 °C to 800 °C with a heating rate of 10 °C $min^{-1}$ under a nitrogen atmosphere.

**Transmission electron microscopy (TEM)**

Transmission electron microscopy measurements were obtained through FEI TECHNAI, USA, FP 5022/22-Tecnai G2 20 S-TWIN transmission electron microscope, which operates at 200 keV using $LaB_6$ filament as an excitation source.

**Fourier transform infrared (FTIR) spectra**

Fourier transform infrared (FTIR) spectra of dried CNDs, were acquired by using a Perkin-Elmer FTIR spectrophotometer equipped with a horizontal attenuated total reflectance (ATR) accessory containing a zinc selenide crystal and operating at 4 $cm^{-1}$ resolution. Spectrum was recorded using Resolutions Pro FTIR software by subtracting background spectra from the sample. The wavelength range to record FTIR spectra of CNDs is 400-4000 $cm^{-1}$.

**Raman spectrum**

The Raman spectrum of dried CNDs was measured by the confocal microscope Raman spectrometer (Horiba Scientific, Xplo RA ONE) with a 532 nm laser (spectral range 400 $cm^{-1}$ to 3500 $cm^{-1}$).

**X-Ray Photoelectron Spectroscopy (XPS)**

Surface chemistry of the material and elemental composition is measured by using XPS in which Auger Electron Spectroscopy (AES) Module PHI 5000Versa Prob II, FEI Inc. and $C_{60}$ sputter gun have been used for the characterization and scanning the spectra for $C_{1S}$, $N_{1S}$, $S_{2p}$, $O_{1S}$ region.

**Confocal Imaging of HeLa Cells**

**Coverslip preparation:** The glass slides and coverslips were cleaned by incubating in freshly prepared Piranha solution for 30 min and finally washing with 70% ethanol in bath sonication, then dried at room temperature.

**Cell Culture, fixation, and staining:** All the cell culture experiments and slide preparations were performed in compliance with the relevant guidelines and norms of biosafety level-1 requirements of the Indian Institute of Technology Mandi. Cell lines were maintained by following the recommended protocols of the National Centre for Cell Science (NCCS) Complex, Pune, Maharashtra, India - 411 007.

Human cervical cancer (HeLa) cell lines and HEK were cultured in Dulbecco's Modified Eagle Medium (DMEM) with 10% fetal bovine serum (FBS), 1% anti-anti, and penicillin/streptomycin at 37 °C with 5% $CO_2$ humidity. The cells were grown in a 6-well plate on coverslips with $10^4$ cells per 100 µl density. Each well was filled with 2 ml of growth medium, and the cells were allowed to grow overnight for proper adherence and growth. Cell growth and attachment to the coverslips were examined with an optical microscope. Once the cells reached the proper adherence and confluency, they were stained with synthesized CNDs fluorescent probe to achieve enough labeling density for confocal microscopy. Finally, the cells were fixed by incubating with 4% paraformaldehyde solution in 1X PBS buffer for 15 min. The fixed cells were washed 2-3 times with PBS buffer to remove extra CNDs and the Cell culture medium. The coverslips were fixed with the help of mounting media (9 glycerol: 1 1X PBS) on a glass slide before imaging.

**ROS detection:** ROS detection was achieved by seeding the cells (40 X $10^3$ cells per well). After the proper adherence of cells, they were incubated with CNDs for 24 h. After the incubation, 1µM of $H_2$DCFDA was added to the cell medium and incubated for 30 minutes. Finally, the fluorescence intensity of cells was measured at 520 nm after excitation with 490 nm, and identification of ROS was achieved and compared with the control sample.

**Cell cytotoxicity:** Cell cytotoxicity was detected with the help of an XTT kit (Roche XTT kit II). HeLa cells were seeded in 96 well plates and cultured overnight with a density of $4 \times 10^3$ cells per well. Then cells were incubated with CNDs for 2 h, at 37 °C in a $CO_2$ incubator. After the incubation with CNDs, the XTT labeling mixture was added and further incubated for 12-16 h. Then, the absorbance of the sample mixture was measured by a microplate reader at 550 nm of wavelength. The reference wavelength was set at 650 nm, and then the cell viability against CNDs was analyzed, comparing it to blank and control samples. The mean and standard deviation were calculated from five wells tested in parallel.

**Confocal microscopy:** Confocal imaging of HeLa cells was performed using Nikon Eclipse Ti inverted microscope, and images were acquired using NIS-Elements software. The cell samples were excited using 560 nm and 639 nm lasers. Finally, the images were collected by choosing a proper filter set.

**SRRF Imaging**

We acquired multi-frame confocal images consisting of 500 frames in a stack, utilizing a 100X oil immersion objective on a Nikon Ti inverted microscope operating in continuous mode during time-lapse. Videos were captured using Nikon's NIS-Elements software using pinhole size:1 AU, scan size:128 x 128 pixels, scan speed: 4 fps, line averaging: disabled, Channel Series: none. The videos were processed for SRRF using the NanoJ-Core and NanoJ-SRRF plugins of ImageJ, which are available at https://sites.imagej.net/NanoJ-SRRF/plugins/ . We developed an ImageJ macro language script to perform batch SRRF analysis on a high-performance GPU-enabled computer (with NVIDIA GeForce RTX 3070). We used default settings with a ring radius of 0.5, a radiality magnification of 5, and axes in ring 6. All videos were first drift-corrected using the ImageJ NanoJ-Core plugin.
